# *De Novo* design of a potent Wnt Surrogate specific for the frizzled7 subtype members

**DOI:** 10.1101/2025.07.28.667009

**Authors:** Junyup Lee, Yeongnam Bae, Tae-Keun Jeong, Seokmin Yun, Jeeyoon Chang, SangPhil Ahn, Hyeongyu Han, Jieun Mun, Seung Jun Kim, Ho Min Kim, Dae-Sik Lim, Yoon Ki Kim, Eui-Jeon Woo, Hanseul Yang, Ho-Chul Shin, Bo-Seong Jeong, Byung-Ha Oh

## Abstract

In humans, 19 Wnt ligands interact with 10 Frizzled (Fzd) receptors and the co-receptors LRP5/6 to initiate signaling. Wnts and Fzds are highly promiscuous, making it challenging to dissect the specific outcomes of individual Wnt-Fzd interactions. Developing Wnt surrogates with specificity for individual Fzd subtypes could be pivotal. We present a modular, potent, and Fzd7-specific Wnt surrogate that consists of three *de novo* designed modules, a Fzd7 binder, an LRP6 binder and a homodimeric protein. The Fzd7-specific module was designed by targeting two less conserved surface patches on the cysteine-rich domain (CRD) of Fzds to achieve both selectivity and affinity. It exhibits a strong binding affinity (*K*_D_ < 2.3 nM) for the very closely related Fzd7 subtype members (Fzd7, Fzd1, Fzd2) with no measurable binding to the CRDs of the other seven Fzd receptors. This Wnt surrogate induced spheroid organoid formation from intestinal stem cells at subnanomolar concentration, and promoted full hair follicle regeneration and robust hair growth in mice. These results suggest that our strategy could be extended to design modular Wnt surrogates capable of selectively activating individual Fzd receptors, providing a valuable tool kit for development and differentiation, organoid cultures and targeted regeneration.

## INTRODUCTION

The human genome encodes 19 Wnt proteins. They are secreted glycoproteins whose signaling pathways can be classified into two main branches: canonical Wnt signaling (β-catenin-dependent pathway) and non-canonical Wnt signaling (β-catenin-independent pathway) ^1, 2^. The canonical pathway, also known as the Wnt/β-catenin signaling pathway, regulates stem cell proliferation and differentiation across a spectrum of biological contexts, encompassing embryonic morphogenesis and adult homeostatic processes. Wnt, the Fzd GPCRs, and its co-receptors LRP5/6 are key components of the Wnt/β-catenin signaling pathway. Wnt initiates this signaling when it forms a ternary complex with Fzd and LRP5 or LRP6, leading to the recruitment of disheveled (Dvl) and Dvl-dependent phosphorylation of the LRP5/6 intracellular domain ^3^. Dysregulation of the Wnt pathway is intricately linked to the pathogenesis of oncological conditions, skeletal anomalies, neurodegeneration, and other pathological states ^4, 5, 6,7^.

The human genome encodes 10 Fzd proteins (named Fzd1 through Fzd10), which belong to class F GPCRs ^8^. Wnts and Fzds are highly promiscuous, and the specific outcomes of interactions between individual Wnt ligands and Fzd receptors remain poorly understood. In a given tissue, Wnt/Fzd/LRP pairing might be influenced by the differential expression of the ligand and coreceptors across different cell types, determining signaling outcomes. Recently, single-cell RNA sequencing revealed that AT2 cells in the human lung highly express Fzd5 and Fzd6, while stromal and endothelial cells predominantly express Fzd4 and Fzd1, respectively ^9^. By leveraging the cell-type-specific Fzd expression, the regeneration of the alveolar epithelium in the mouse lung was possible through selective activation of Wnt signaling using Fzd-specific agonists ^10^, highlighting the potential to disentangle the complexity of Wnt-Fzd signaling pathways involved in stem cell activation by combined use of such agonists.

In research applications, common methods for activating Wnt signaling, such as using recombinant Wnt3a or small molecular GSK3 inhibitors, effectively trigger downstream signaling but lack receptor specificity, limiting their usefulness in studying biological responses of specific Wnt-receptor interactions ^11^. Recombinant Wnt3a, widely used as a Wnt source for organoid culture, is lipidated and requires conditioned media for solubility ^12^, which can introduce variability and negatively impact certain organoid types ^13^. Additionally, GSK3 inhibitors like CHIR broadly activate Wnt signaling but can also affect other cellular pathways, potentially causing unintended side effects ^14^. These issues underscore the need for receptor-specific Wnt surrogates to achieve precise and controlled activation of Wnt signaling. To address these limitations, water-soluble Wnt surrogates have been developed based on a bacterial four-helical bundle or ankyrin repeat scaffolds ^15, 16^ or antibodies which bind the CRDs of multiple Fzds ^17, 18^, and the C-terminal fragment of Dickkopf-related protein 1 which binds to LRP5/6. While these surrogates represent a significant contribution to organoid research, availability of individual Fzd-specific Wnt surrogates would be valuable tools for further advancing both therapeutic and research applications. Designing or engineering such proteins is challenging due to the high sequence homology among the 10 human Fzd proteins.

Here, we present the *de novo* design of a protein that binds to the CRD of human Fzd7/Fzd1/Fzd2 with high affinity, without cross-reactivity to those of the other 7 human Fzd subtypes. In addition, we *de novo* designed LRP6-binding protein and a homodimeric coiled-coil protein. By fusing the Fzd7-specific binder with the LRP6-binder, we created a Wnt surrogate. When further dimerized using the designed coiled-coil protein, this fusion protein forms a potent Wnt agonist.

## RESULTS

### *De novo* design of Fzd7-specific binder

The Fzd receptors have a small ectodomain composed of about 110 residues, called cysteine-rich domain (CRD). Wnt binds to the CRD of Fzd with two specific binding sites resembling ‘thumb’ and ‘index finger’, respectively ^19^. Wnt’s thumb is modified with a palmitic acid, and this lipid moiety is the major motif interacting with the hydrophobic cleft of the CRD (Figure 1A and 1B). On the other hand, Wnt’s index finger interacts with a concave surface on the opposite side of the CRD. To design binder protein that preferentially interacts with Fzd7, we utilized the ConSurf server ^20^ to assess the conservation of surface amino acids of the CRD domains across 10 human Fzd receptors (Figure 1A). The lipid-binding site is highly conserved and hydrophobic, whereas the Wnt’s index finger-interaction surface is more variable and hydrophobic (Figure 1A, B). We selected two surfaces for binder design: a variable patch adjacent to the lipid-binding groove (Site 1) and the Wnt’s index finger-interaction surface (Site 2) (Figure 1A, B; boxed regions). Due to their opposing locations, we opted to design two separate proteins, each targeting one of the surfaces. For Site 1, we identified and designated three hydrophobic residues (Leu81, Pro88, Phe138) on this surface as key interacting residues (hot spots), and generated protein backbones using RFdiffusion ^21^. The sequences of these backbones were then designed using ProteinMPNN ^22^. Semi-validation based on the structure prediction using AlphaFold-Multimer (AF-Multi) ^23^ was performed to select final candidates for experimental validation (Figure 1C). A total of 62 designed proteins were produced, and their binding to the CRD of Fzd7 was assessed using biolayer interferometry (BLI). One of the designed proteins was outstanding in the binding kinetics, and its dissociation constant (*K*_D_) was quantitated to be 238.2 nM. While promising, the binding affinity is substantially low compared with a measured affinity of Wnt (*K*_D_ of 2.8 nM) ^24^. We sought to extend the C-terminus of this Site 1 binder to form an additional protein domain that interacts with Site 2. This second design was based on the AF2Multi-predicted structure of the Site 1 binder bound to Fzd7 CRD and on using a similar design protocol (Figure 1D). For each design a linker between the two domains were also designed. Four design candidates were selected for protein production and experimental validation by quantification of the binding affinity for Fzd7 CRD. The finally selected design, having a short linker between the Site 1 binder and the Site 2 binder, is reminiscent of a headset (Figure 2A). This design was named Wnt_M1 (for Wnt mimetic 1).

**Figure 1.**
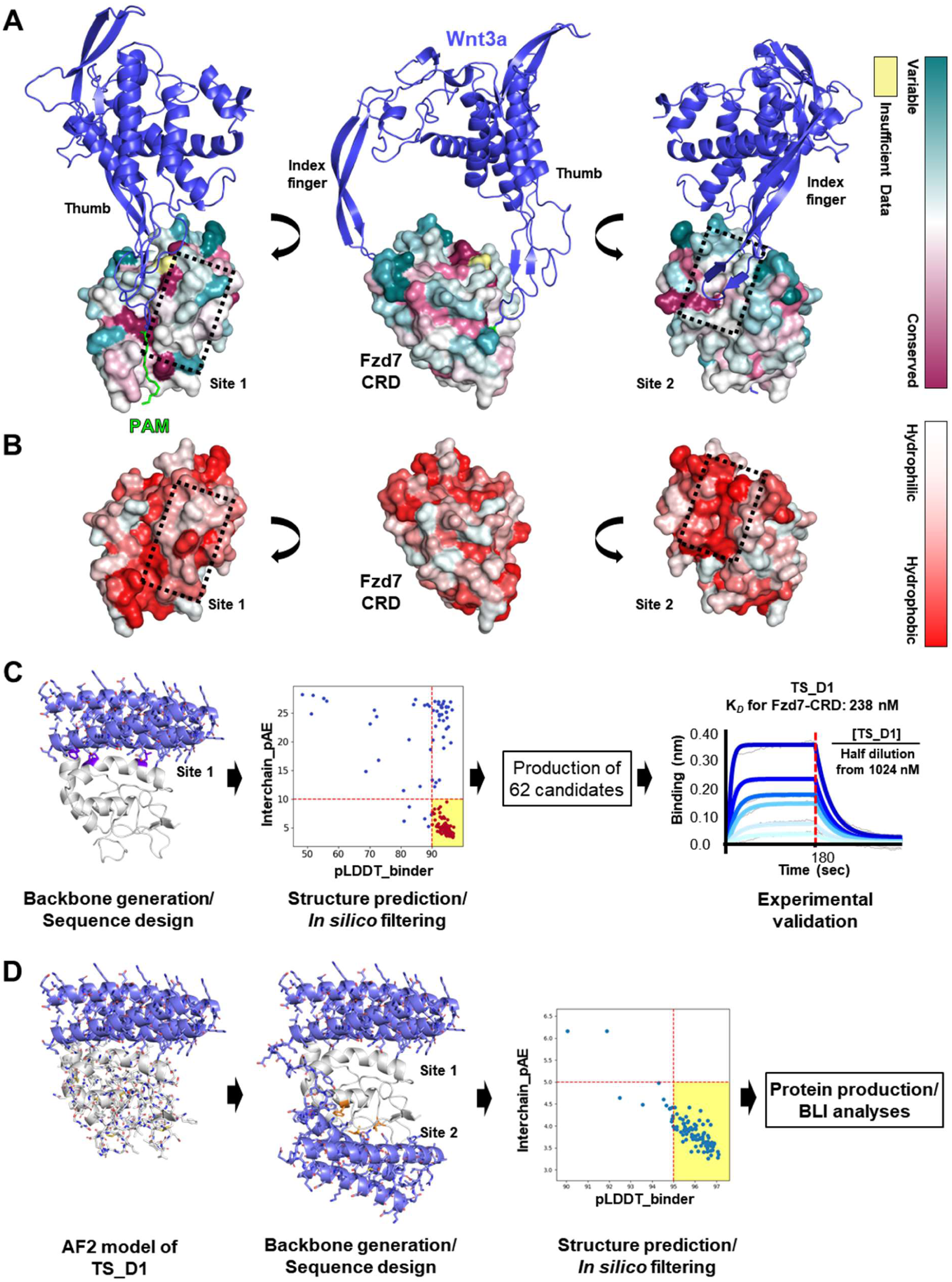
Selection of target surfaces and design protocol for Fzd7-Specific binders. **A.** Surface conservation. The structure of human Wnt3a bound to the cysteine-rich domain (CRD) of Fzd7 was predicted using AF-Multi and used to illustrate the two distinct binding interfaces. The palmitoleic acid on the ‘Thumb’, shown in green, was adapted from the crystal structure of Xenopus Wnt8 bound to the CRD of mouse Fzd8 (PDB entry: 4F08). The CRD is color-coded to illustrate the surface residue conservation among the 10 human Fzd proteins. **B.** Hydrophobic surface representation. The two interaction sites on the CRD are notably hydrophobic. The dotted boxes indicate the two selected target surfaces for the *de novo* binder design. **C.** Computational procedure for Site 1 binder design. The hot-spot residues for the backbone generation are shown in purple. The structures of the designs were predicted using AF-Multi, and those with scores highlighted in the yellow box were selected for experimental validation. **D.** Computational procedure for Wnt_M1 design. Backbone generation was guided by designated hot-spot residues, shown in orange. Among the designs with scores highlighted in the yellow box, four were selected for experimental validation based on visual inspection.

**Figure 2.**
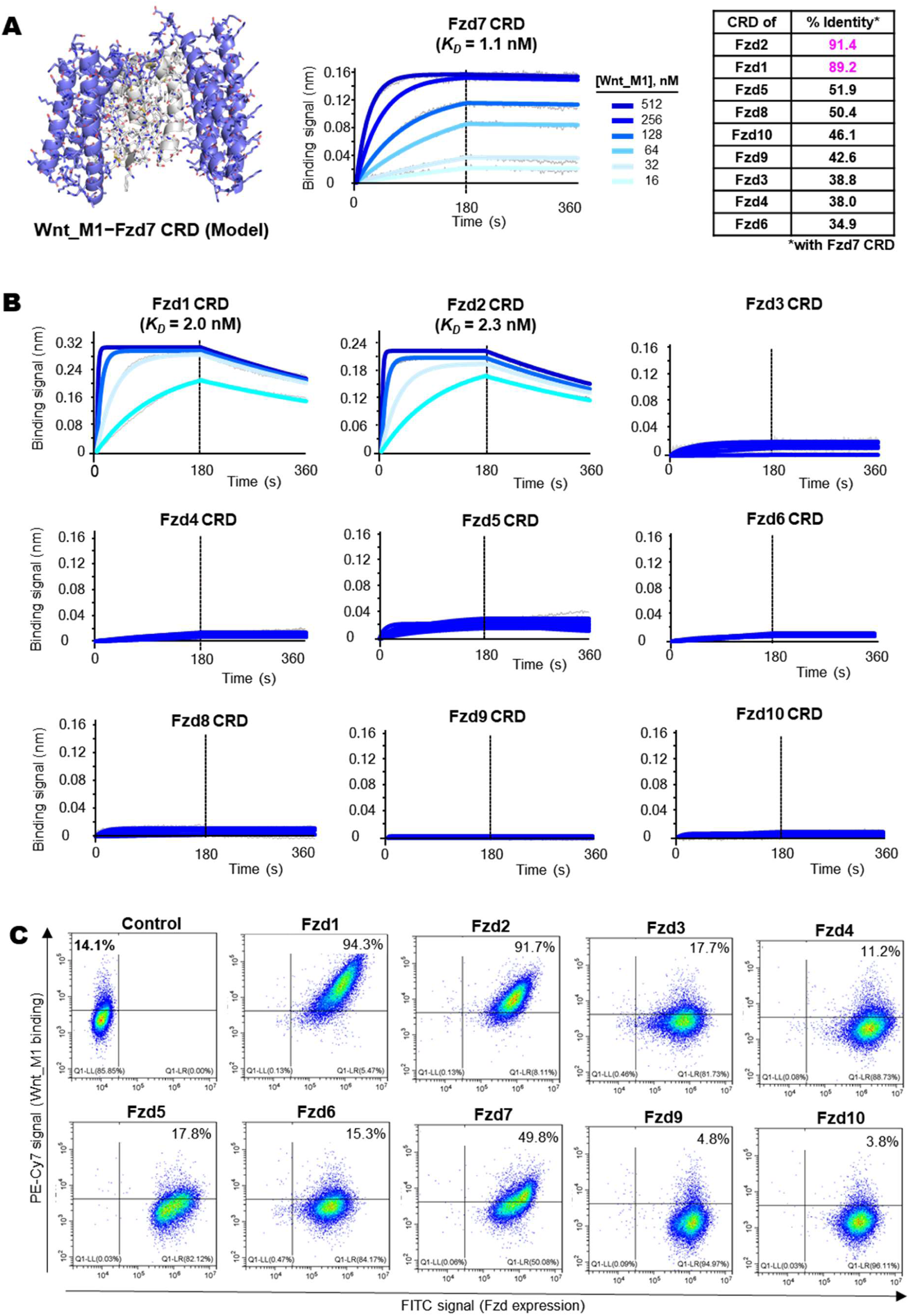
Binding affinity and specificity of Wnt_M1. **A.** Structural model of Wnt_M1 (*Left*) and quantification of the binding affinity for Fzd7 CRD (*Right*). Biotinylated Fzd7 CRD at 400 nM was immobilized on SA-tips and incubated with Wnt_M1 at the indicated concentration. The *R*^2^ value for curve fitting was 0.9871. Pairwise sequence identities between the Fzd7 CRD and those of other subtypes are tabulated. **B.** Binding affinity for other human Fzd subtypes. Biotinylated target at 400 nM was captured on SA-tips and incubated with Wnt_M1 at concentrations from 1024 nM to 32 nM in a serial two-fold dilution. Binding affinities were measured for the CRDs of Fzd1 and Fzd2. Binding affinity was estimated using the Octet ForteBio software, with curve fitting performed using a 1:1 steady state model. The *R*^2^ values for curve fitting were 0.9923 and 0.9971 for Fzd1 CRD and Fzd2 CRD, respectively. For the CRDs of other Fzd subtypes, binding was too weak to determine affinities. **C.** Binding specificity of Wnt_M1. ExpiCHO cells expressing exogeneous full-length Fzd receptors were incubated with Myc-tagged Wnt_M1 (100 nM). Bound ligand was detected by flow cytometry using a phycoerythrin-conjugated anti-Myc antibody. In the control panel, 14.1% of untransduced ExpiCHO cells exhibited background-level binding signals above the threshold. The relatively weaker signal from Fzd7-expressing cells, in comparison with Fzd1- or Fzd2- expressing cells, is likely due to instability of Fzd7. Attempts to generate Fzd8-expressing cells were unsuccessful owing to cloning challenges posed by extensive glycine repeats.

### Binding affinity and specificity of Wnt_M1

As expected from the avidity effect, Wnt_M1 exhibited a substantially enhanced binding affinity for Fzd7 CRD, with a *K*_D_ of 1.12 nM, approximately 210 times higher than the Site 1 binder (Figure 2A). To evaluate the binding specificity of Wnt_M1, BLI was performed using the CRDs of the other nine human Fzd receptors. Wnt_M1 showed high-affinity binding to Fzd1 and Fzd2, both of which share over 89% sequence identity with the Fzd7 CRD, but exhibited no noticeable binding to the CRDs of the remaining seven Fzd subtype members (Figure 2B), demonstrating exceptional selectivity for the Fzd7 subtype members. To determine whether this binding specificity is preserved in a cellular context, full-length Fzd receptors were expressed on Expi-Chinese hamster ovary (CHO) cells, and binding to Wnt_M1 was analyzed by flow cytometry. Consistent with the BLI results, Wnt_M1 exhibited strong binding to cells expressing Fzd1, Fzd2 or Fzd7 (Figure 2C). Cells expressing the other Fzd receptors did not show appreciable binding to Wnt_M1 above the control level. These results indicate that Wnt_M1 can effectively distinguish the Fzd7 subtype members from other Fzd subtypes on cells, and validates our approach of targeting two less conserved surfaces to achieve both exceptional selectivity and tight binding affinity.

For comparison, we performed the same cell-binding analysis with the previously reported Fzd7-specific single-domain binder DRPB_Fz7, which was designed to target both the highly conserved fatty acid-binding groove and an adjacent, less conserved surface patch ^25^. In addition to Fzd1-, Fzd2-, Fzd7-expressing cells, considerable binding signals from Fzd5-, Fzd6-, Fzd10-expressing cells were observed. Subsequent BLI analysis at multiple concentrations showed that DRPB_Fz7 binds to the Fzd5 CRD with a 168.5 nM (Supplementary Figure 1).

### *De novo* design of LRP6 binder

To create a Wnt-mimetic protein that can initiate Wnt/β-catenin signaling through simultaneous binding to Fzd7 and LRP6, we designed a *de novo* LRP6 binder for fusion with Wnt-M1. LRP6 is a single-pass transmembrane protein, whose extracellular portion is composed of four similar domains, designated as E1 through E4 from the N-terminus to the C-terminus, each composed of YWTD-type β-propeller domain and an epidermal growth factor-like domain (Figure 3A). We utilized the structure of the complex between the LRP6 E1 domain and antigen-binding fragment (Fab) of the antibody YW210.09 ^26^. The key 6-amino acid paratope (VNAVKN) of the antibody contains the core binding motif (NAV), which aligns with the conserved NXV motif, where X is Ala, Ser or Trp, found in other known LRP6-binding proteins ^26^. The paratope structure was used as a binding motif in generating various backbone structures incorporating this motif using RFdiffusion. Amino acid sequences were then designed with ProteinMPNN, and the complex structures between each design and the target were predicted using AF-Multi (Figure 3B). From models with pLDDT > 90 and interchain pAE < 10, eight that preserved the binding motif conformation were selected for experimental validation by measuring their binding affinity for biotinylated LRP6 E1-E2 (Figure 3C). The best design, exhibiting the highest binding affinity of (*K*_D_= 12.9 nM), was finally selected and named LRP6 binder (Figure 3D). We presume that this LRP6 binder would also binds to the LRP5 E1 domain, because the E1 domains of LRP5 and LRP6 share 73 % sequence identity with each other, and the epitope residues interacting with the 6-amino acid paratope are identical between the two LRP proteins.

**Figure 3.**
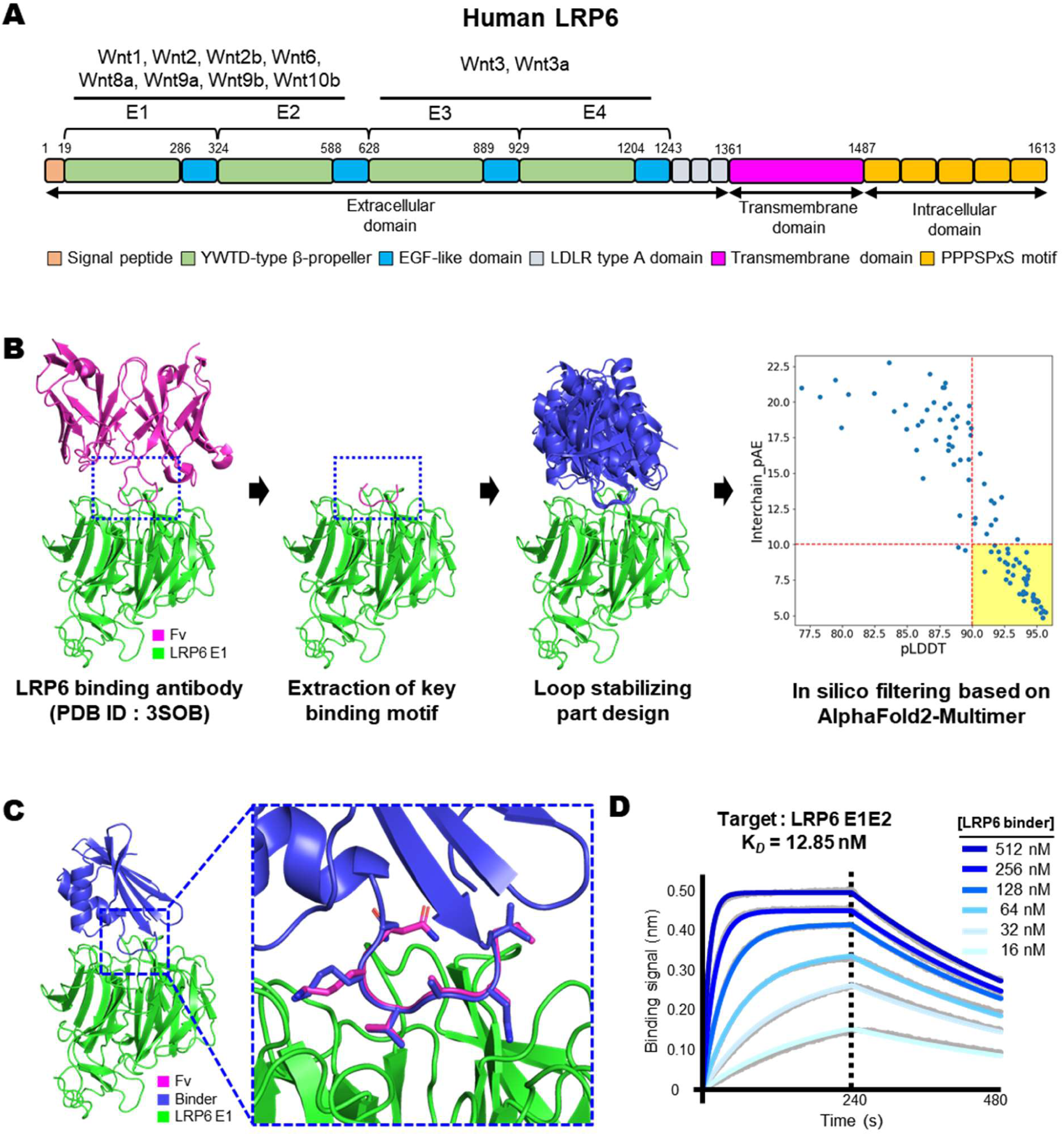
*De novo* design of LRP6 binder. **A.** Schematic diagram of human LRP6. Known interactions between Wnt proteins and the E1-E2 or the E3-E4 domain are indicated. **B.** Computational procedure. The key binding motif was extracted from a LRP6 E1E2-binding antibody and retained in the backbone and sequence designs. The designed sequence was assessed by AF-Multi. Of the designs with scores in the yellow box, eight designs were selected for experimental validation based on visual inspection. **C.** An example of structural model that preserved the conformation of the input binding motif (magenta). **D.** Binding affinity of the finally selected LRP-6 binder. BLI was performed by capturing 400 nM biotinylated LRP6 E1E2 on SA-tips, incubating with the LRP6 binder at the specified concentrations, and proceeding through the dissociation phase. Curve fitting was performed using a 1:1 steady state model.

### *De novo* design of a homodimer-forming protein module

The LRP6-binding module was fused to Wnt_M1 via a 7-residue linker (GGGSGGG), generating a fusion protein capable of binding both Fzd7 and LRP6. To enhance binding avidity and thereby increase Wnt signaling potency, we aimed to dimerize this fusion protein using an elongated homodimerization module. For this purpose, we employed the ColE1 repressor of primer (Rop) ^27^. We extracted the central region of the Rop coiled coil and symmetrically extended its helices at both termini using RFdiffusion. The sequences of the newly generated helical extensions were designed with ProteinMPNN, and candidate designs were selected based on AlphaFold2 (AF2) confidence scores (Figure 4A). Among the 12 top-scoring designs, 10 were expressed in soluble form. The best-performing construct, named HomoCC, was chosen for its favorable solution behavior (Figure 4B, Left). The crystal structure of HomoCC was determined at 2.0 Å resolution and closely matched the design model (Figure 4B, Right).

**Figure 4.**
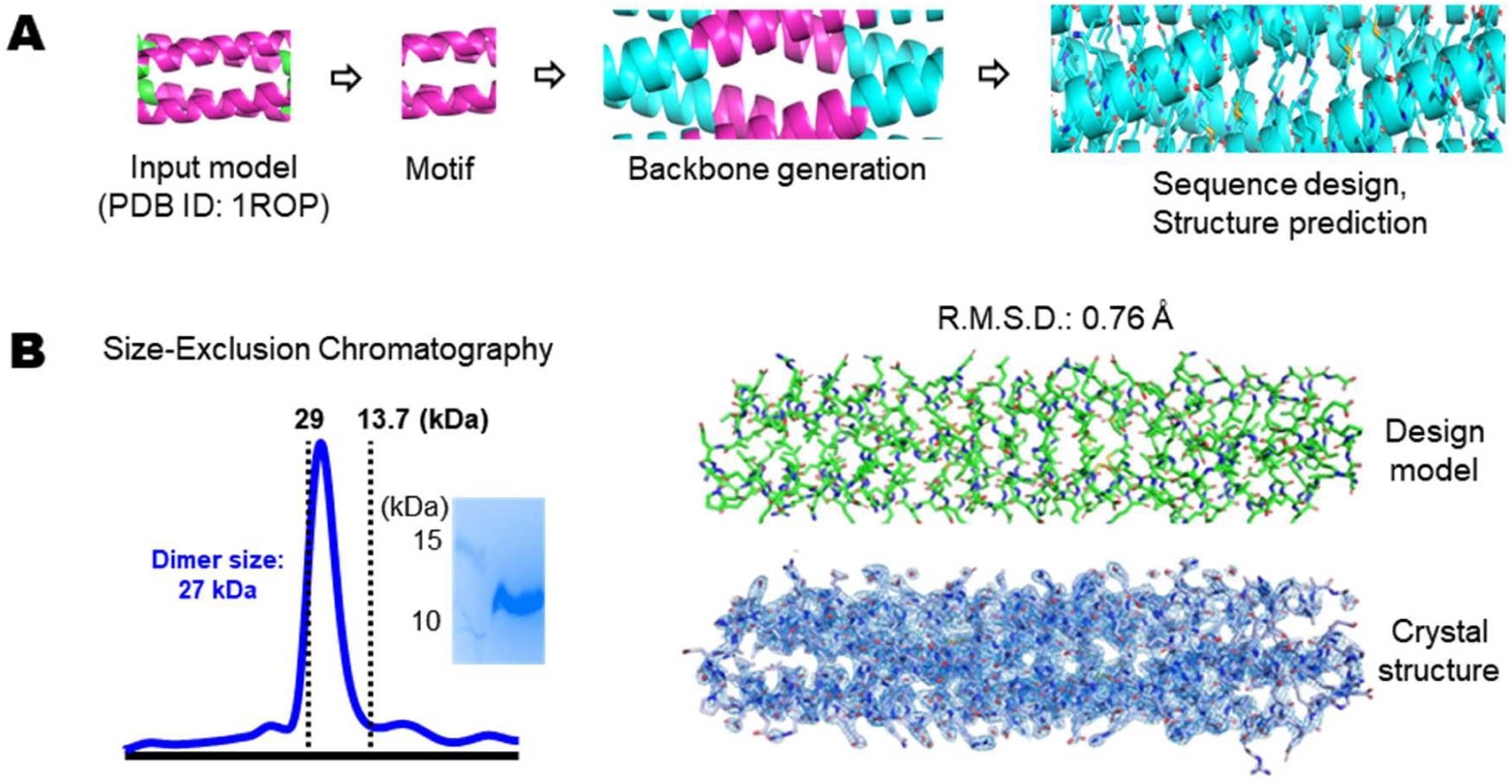
Computational design of HomoCC and generation of modular Wnt surrogates. **A.** Computational procedure. In the antiparallel coiled coil (PDB entry: 1ROP), the core portion (in magenta) was used as a motif from which α-helical backbones were extended and their sequences were designed by RFdiffusion and ProteinMPNN, respectively. **B.** Solution behavior and structure determination of HomoCC. The protein was eluted as a single peak and as if it is a homodimer from a size-exclusion chromatographic column. This fraction showed a single peak on a SDS-PAGE gel (*Left*). The crystal structure of HomoCC and the predicted design model are shown in the same orientation. The superposition of the two structures results in an R.M.S.D. of 0.76 Å for all aligned Cα atoms (*Right*).

### Generation of modular Wnt surrogates and their agonistic activities

To generate a modular Wnt surrogate capable of binding both Fzd7 and LRP6, the LRP6-binding module was fused to Wnt_M1 via a 7-residue flexible linker (GGGSGGG), yielding a monomeric fusion protein termed F7_WS_mono (Fzd7-specific Wnt surrogate monomer). To enhance binding avidity and thus signaling potency, F7_WS_mono was further fused via a 5-residue linker (GGSGG) to HomoCC, forming a homodimeric construct F7_WS_di (Fzd7-specific Wnt surrogate dimer). This dimeric format would enable simultaneous engagement of two Fzd7-LRP6 receptor pairs and thus receptor clustering (Figure 5A). Both F7_WS_mono and F7_WS_di were highly expressed in soluble form in *E. coli* RIPL cells and purified to a highly homogeneous state (Figure 5B).

**Figure 5.**
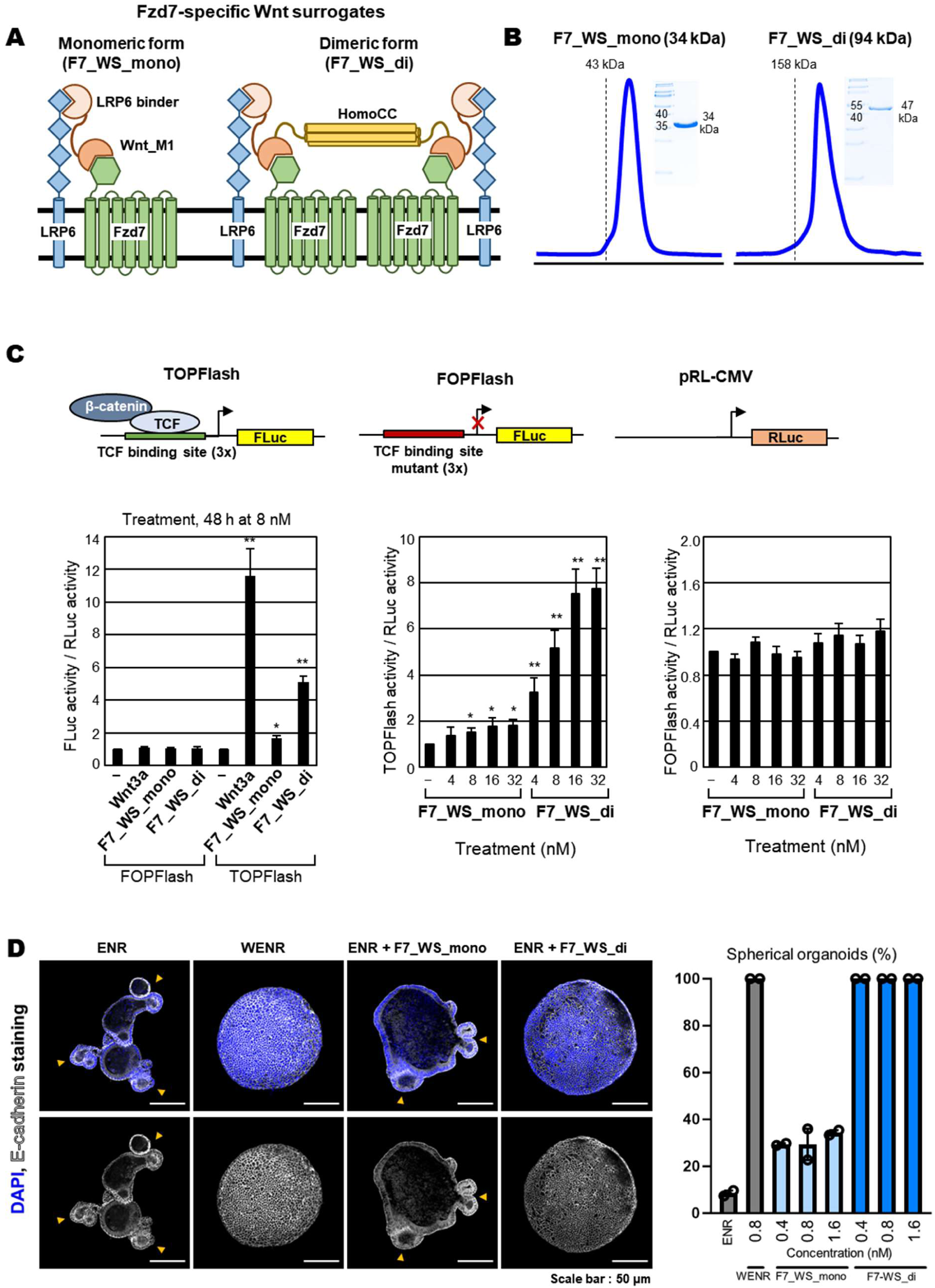
Modular Wnt surrogates function as agonists of Wnt/β-catenin signaling. **A.** Schematic drawing of the fusion protein binding to Fzd7 and LRP6; The dimeric version includes HomoCC for self-dimerization. **B.** Size-exclusion chromatography and SDS-PAGE show the homogeneity of the monomer (F7_WS_mono) and dimer (F7_WS_di) constructs. **C.** TOP/FOPFlash reporter assay for Wnt/β-catenin signaling in HCT116 cells. *Top*: reporter schematics. Wnt3a is an activator of TCF-dependent transcription. *Bottom, Left*: Cells were transiently transfected with either TOPFlash or FOPFlash and a reference plasmid expressing Renilla luciferase (RLuc). 8 nM Wnt3a or F7_WS_di, but not F7_WS_mono, markedly activates TOPFlash. The normalized FLuc levels in untreated cells were set to 1.0. *Bottom, Middle*: F7_WS_di elicits a far stronger, dose-dependent TOPFlash response than F7_WS_mono. *Bottom right*: FOPFlash activity remains at baseline. Data, mean ± SD (n = 3); *P < 0.05, **P < 0.01. **D.** Murine intestinal organoid culture. *Left*: confocal images after 3 d in ENR medium containing 1.6 nM WENR (Fc-Wnt surrogate), F7_WS_mono, or F7_WS_di. E-cadherin (white) outlines epithelia; DAPI (blue) labels nuclei. In the F7_WS_mono group, some organoids exhibited budding forms, indicated by arrowheads. *Right*: Quantification of spherical organoids after exposure to 0.4-1.6 nM surrogates. Whereas F7_WS_mono yielded fewer than 40 % spherical organoids, F7_WS_di generated a uniformly spherical organoid population.

To assess their signaling function *in vitro*, we used the TOPFlash and FOPFlash luciferase reporter systems incorporated into HCT116 cells, which express Fzd7 ^28^. These constructs contain the firefly luciferase (FLuc) gene driven by a thymidine kinase minimal promoter with either wild-type (TOPFlash) or mutated (FOPFlash) TCF-binding sites ^29^ (Figure 1A). In cells co-transfected with TOP/FOPFlash and a Renilla luciferase control, Wnt3a, an established activator of TCF-dependent transcription ^29^, selectively increased TOPFlash activity, validating the assay (Figure 5C). Under identical conditions, the dimeric F7_WS_di, but not the monomeric F7_WS_mono, significantly induced TOPFlash activity (Figure 5C). Much greater dose-dependent stimulation of the TOPFlash activity by F7_WS_di confirms the greater signaling potency of the dimer over the monomeric form (Figure 5C). These results demonstrate that the *de novo* designed dimeric modular protein functions as an effective Wnt surrogate, capable of activating the canonical Wnt/β-catenin signaling.

We next tested the potency of the Wnt surrogate in generating organoids from mouse Lgr5^+^ intestinal stem cells, where Fzd7 is known to be a key subtype in mediating Wnt signaling ^30^. Murine intestinal organoids exhibit distinct morphologies depending on the presence of Wnt ligands. In an ENR medium lacking a Wnt, organoids generally grow in a budding form with crypt-like domains. In contrast, in a WENR medium containing a Wnt ligand, they adopt a spherical morphology. Consistently, mouse stem cells cultured in ENR and WENR media predominantly grew in budding and spherical forms, respectively. Upon treatment of F7_WS_mono, organoids exhibited an increased number of spherical forms, but still contained budding forms as well. However, with treatment of F7_WS_di, all organoids adopted a spherical form, similarly observed with the WENR treatment (Figure 5D). Notably, this effect was attained at concentrations lower than 0.8 nM, an effective concentration of a commercial WNT surrogate-Fc fusion protein, which is also a homodimeric form of a *de novo* designed Fzd-binder fused to the Dickkopf 1 C-terminal domain ^16^. These results confirm that both F7_WS_mono and F7_WS_di function as a Wnt ligand in the murine intestinal organoid model, and the homodimeric form is far more potent as expected from the increased avidity effect. Furthermore, F7_WS_di provides the optimal 2:2 stoichiometry of Fzd7 and LRP binders, aligning with prior work showing that tetravalent binding to Fzds and LRPs is a requirement for maximal WNT/β-catenin activation ^18^.

### *In vivo* hair follicle regeneration and hair growth by F7_WS_di

Recently, the role of Wnt7a in hair follicle regeneration was demonstrated in a wound-induced hair neogenesis model using *K14-Wnt7a* transgenic mice, where Wnt signal amplification was achieved through epidermal overexpression of Wnt7a ^31^. Given that hair-follicle stem cells (HFSCs) express high levels of Fzd2 and Fzd7, and Wnt ligands initiate hair follicle regeneration cycle ^32^, we tested whether our Wnt surrogate could activate HFSCs. F7_WS_di was injected intradermally on postnatal days 60 and 62, corresponding to the telogen phase when HFSCs are in a resting phase (Figure 6A). Three days after the second injection, F7_WS_di at 1 µM and 4 µM concentrations induced robust HFSC proliferation, as evidenced by 5-Ethynyl-2′-deoxyuridine (EdU) incorporation and co-localization with the Wnt/β-catenin target LEF1 (Figure 6B,C). By day 14, the same treatment led to fully regenerated hair follicles and visible hair growth (Figure 6D,E). These results demonstrate that the Fzd7-specific Wnt surrogate effectively activates canonical Wnt/β-catenin signaling *in vivo*, driving HFSC activation and complete hair-follicle regeneration. This treatment, corresponding to 10 or 40 μg protein injection under the skin did not accompany any sign of adverse effect (Figure 6E). The hair follicle regeneration and hair growth likely require long-lasting agonistic effect of F7_WS_di. Circular dichroism (CD) spectroscopic analyses support protein stability of F7_WS_di; Wnt_M1 and HomoCC showed no apparent unfolding transition up to 90 °C, while LRP6 binder showed unfolding transition above 80 °C (Supplementary Figure 2).

**Figure 6.**
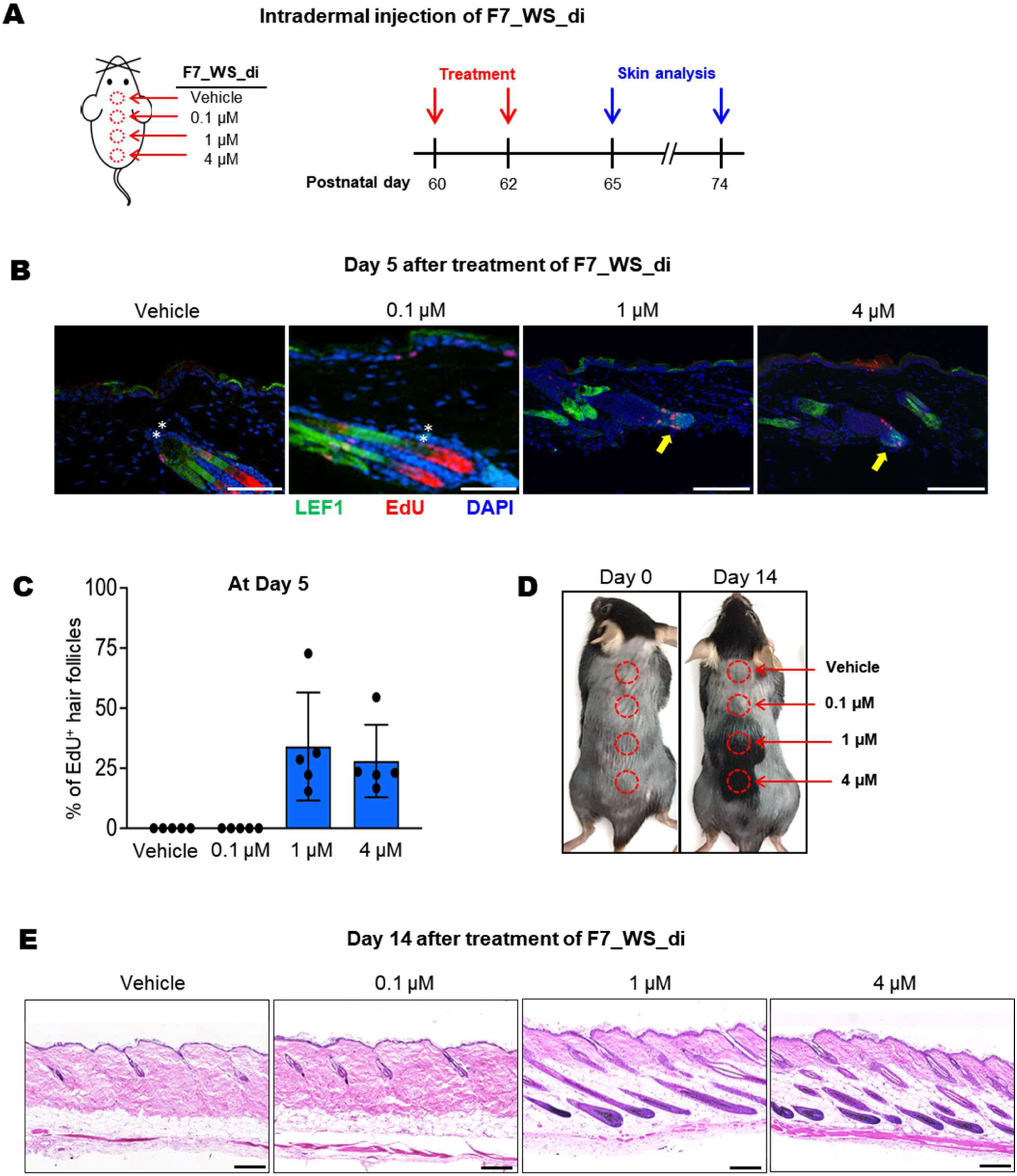
*In vivo* hair follicle regeneration induced by F7_WS_di. **A.** Schematics of the experimental design. **B.** Immunofluorescence images showing LEF1 expression (a readout of Wnt/β-catenin signaling) and EdU incorporation (a marker of cell-cycle entry) in hair follicle stem cells. Yellow arrows indicate proliferating EdU⁺ HFSCs. Scale bars, 100 μm. Asterisks denote autofluorescence signals from hair shafts. **C.** Quantification of EdU⁺ hair follicles (n = 5 mice). *p < 0.01; **p < 0.001. **D.** Intradermal injection of 50 μL of F7_WS_di (1 μM and 4 μM) induced robust hair growth. **E.** Hematoxylin and eosin staining showing fully regenerated hair follicles in skin treated with 50 μL of F7_WS_di at concentrations of 1 μM and 4 μM. Scale bars, 100 μm. The tissue structure remained intact.

## DISCUSSION

In many ligand-receptor signaling pathways, multiple receptor subtypes with high sequence homology interact with ligands, which also share homologous sequences, to produce diverse and pleiotropic signaling outcomes. A typical example is the Wnt-Fzd-LRP5/6 signaling pathway, involving 19 Wnts, 10 Fzds, 2 LRPs. With the large number of Fzd and Wnt subtypes, and their high degree of subtype cross-reactivity, the functional dissection of individual Wnt-Fzd interactions remains a challenging problem. Considerable effort has been put forth to achieve selective activation by developing Fzd subtype-specific antibodies. However, the small size and high sequence homology of the CRDs among Fzd receptors have hindered the discovery of truly subtype-specific antibodies, and most reported antibodies show cross-reactivity with multiple CRDs. One exception is a Fzd7-specific antibody that was developed to bind the linker sequence (the extracellular ‘neck’ region) between the transmembrane domain and the CRD of Fzd ^33^. While this antibody has not been developed as a Wnt mimetic, it demonstrated potent efficacy as an antibody-drug conjugate, killing ovarian cancer cells in vitro and inducing tumor regression in preclinical ovarian cancer models ^34^.

High-affinity, subtype-specific Wnt surrogates based on ankyrin scaffolds were designed to engage the highly conserved fatty acid-binding groove of Fzd CRDs for maximal affinity while simultaneously contacting an adjacent, less conserved surface to impart specificity. Nevertheless, because the lipid-binding groove is conserved across Fzd subtypes, a degree of cross-reactivity beyond the intended subtype is unavoidable (Supplementary Fig. 1).

By simultaneously engaging two less-conserved surface patches on the Fzd7 CRD, rather than a single site, we aimed to create binders with near all-or-none specificity while preserving high affinity. The resulting design, Wnt_M1, binds Fzd7/Fzd1/Fzd2 with a single-digit nanomolar affinity and shows no measurable affinity for the remaining seven Fzd subtypes. With this exceptional selectivity, Wnt_M1 might potentially serve as a Wnt antagonist that could block Fzd7/Fzd1/Fzd2-mediated Wnt/β-catenin signaling with minimal off-target activity. Alternatively, Wnt_M1 could be repurposed as a biosensor to profile Fzd7/Fzd1/Fzd2 expression across cell types—in combination with gene expression analyses—thereby enabling precise spatial and temporal mapping of these subtypes within specific tissues.

Commercially available Wnt agonists typically rely on fusion proteins combining Fzd and LRP binding domains, and they are produced from mammalian cells. By contrast, F7_WS_di is readily produced in bacteria, remains highly soluble, and can be concentrated to at least 300 µM without precipitation. Its exceptional solubility and stability make F7_WS_di a powerful reagent for dissecting the role of the Wnt/β-catenin signaling through the Fzd7 subtype members across diverse cell types, potentially uncovering cell-type-specific roles in tissue regeneration and cancer progression.

The presented Fzd7-specific Wnt surrogate is a modular protein, and the design approach described here offers a promising strategy for designing subtype-specific Wnt mimetics tailored to other Fzd receptor. By replacing Wnt_M1 with protein modules selective for different Fzd subtypes, additional subtype-specific Wnt surrogates could be generated. This modular framework would expand the toolkit for dissecting the roles of individual Fzd receptors across diverse cell types and tissues, in both normal and disease contexts.

In conclusion, this study serves as a showcase, demonstrating that targeting two less conserved surface patches of the CRD domains of Fzd receptors enables the generation of fully *de novo* subtype-specific Wnt mimetic proteins with a headset shape, and that a potent Wnt surrogates with specificity for individual Fzd subtypes can be generated by incorporating this protein as a specificity-conferring module within the Wnt mimetic-LRP binder-HomoCC format.

## METHODS

### *De novo* design of Fzd7-binding protein

To design Site 1 binder, Leu81, Pro88 and Phe138 were designated as key interaction residues (hotspots). Protein backbones consisting of 90-110 amino acids were generated using RFdiffusion ^21^, and compact backbone structures were selected based on a radius of gyration (ROG) less than 15 Å. Amino acid sequence design for the selected models was performed using ProteinMPNN ^22^, generating 100 sequences per backbone. *In silico* prediction using AF2-Multi ^23, 35^ was conducted to assess whether the designed sequences would fold as designed and bind to the target.

To design Site 2 binder, Phe100, Val109 and Val161 were designated as hotspots. Other design steps were similar to those for Site 1 binder. For constructing the connecting loop between the two binders, Val90 and Val92 were selected as hotspots. RFdiffusion was used to generate peptide linkers of 9-12 amino acids in length. Sequence design was then performed on these linkers using ProteinMPNN, and the full-length constructs were evaluated using AF2-Multimer to select candidates for experimental validation.

### *De novo* design of LRP6-binding protein

From the structure of the Fab-LRP6 E1 domain complex (PDB entry: 3SOB), the binding epitope peptide (VNAVKN) was extracted. Using the LRP6 E1 structure, backbone structures of 80-100 amino acids that stabilized the peptide were generated with RFdiffusion. Compact backbones were selected based on a ROG < 13 Å. Amino acid sequence design was performed using ProteinMPNN, generating 50 sequences per backbone. *In silico* structure prediction of protein complexes was carried out using AF2-Multi. For both Fzd7- and LRP6-binding proteins, design models with pLDDT > 90 and interchain pAE < 10 were selected for production and experimental validation.

### *De novo* design of HomoCC

From the crystal structure of ColE1 Rop (PDB ID: 1ROP), coordinates for residues 11–24 and 36– 49 were extracted and used as an antiparallel homodimeric coiled-coil motif. Using RFdiffusion, extended C2 symmetric backbones were created, forming an antiparallel homodimeric coiled-coil structure of 110–120 amino acids. Symmetric sequence design for the generated backbones was carried out using ProteinMPNN, followed by structural evaluation with AF2. Models predicted to form antiparallel dimers with pLDDT > 85 and pTM values > 0.8 were selected for production and experimental validation.

### Expression and purification of the designed proteins

Synthetic DNA fragments encoding the designed proteins were purchased from TwistBioscience, and cloned into a modified version of pET30a vector (Novagen) to produce the protein N-terminally fused to 10x(His)-muGFP-TEV protease cleavage sequence. The proteins were expressed in the *E. coli* BL21 (DE3) RIPL strain at 37 ℃ for 6 h after induction by 0.2 mM IPTG. Cells were harvested and sonicated in Buffer A containing 20 mM Tris-HCl (pH 7.5) and 150 mM NaCl. The supernatant was loaded on to Ni-NTA resin (Thermo Scientific), washed with Buffer A containing additional 20 mM imidazole, and bound proteins were eluted with Buffer A containing additional 250 mM imidazole. The eluted fraction was loaded onto a HiLoad 26/60 Superdex 75 gel-filtration column (Cytiva) equilibrated with Buffer A. Fractions containing the designed protein were pooled and incubated with 10x(His)-tagged TEV protease at 4 °C for 18 h to cleave the 10x(His)-muGFP tag. The reaction mixture was then loaded to Ni-NTA resin, and the flow-through fraction was further purified using the same size-exclusion chromatography. F7_WS_mono, F7_WS_di, Myc-tagged Wnt_M1 and DRPB_Fz7 were produced and purified using virtually the same protocols.

### Crystallization and Structural Determination of HomoCC

HomoCC (20 mg/ml) was crystallized in a solution containing 150 mM MgCl₂, 100 mM Tris (pH 8.5), and 4 M 1,6-hexanediol under sitting drop conditions at 20 °C. Diffraction data were collected at beamline 5C of the Pohang Accelerator Laboratory and processed using the HKL2000 suite ^36^. Phases were determined by molecular replacement with PHASER ^37^, using the AF2-predicted model of HomoCC. Model building and structural refinement were performed with COOT ^38^, PHENIX ^39^ and CCP4 ^40^. Crystallographic data statistics are summarized in Supplementary Table 1.

### Preparation of the CRDs of human Fzd1 to Fzd10

The amino acid sequence of the Fzd7 CRD was used to define the boundaries of the CRDs of the other 9 Fzds by sequence alignment. The DNA fragment encoding each CRD sequence was cloned into a modified pX vector ^41^, which contained the bovine prolactin signal sequence at the 5’ end, and the Precision cleavage site, mCitrine, 10x(His) and biotin acceptor peptide sequence at the 3’ end of the CRD sequence. Expi293F cells (Thermo Fisher Scientific) were transfected with each cloned vector and grown for 10 days at 37 °C. Cell culture media were collected and loaded onto a HisTrap column (Cytiva), which subsequently washed with a buffer solution (150 mM NaCl, 20 mM Tris (pH 7.5), 0.2 mM TCEP), and the bound proteins were eluted with an imidazole gradient (0–0.5 M). The eluted fraction was further purified using a HiLoad 26/600 Superdex 75 pg column (Cytiva) equilibrated with the same buffer.

### Quantification of binding affinity

Binding affinity was measured using an Octet R8 system (Satorius). Biotinylated 6x(His)-mCitrine-tagged CRDs of human Fzd proteins (Fzd1-Fzd10) at 400 nM were immobilized onto streptavidin biosensor tips (Satorius) in Kinetics Buffer (Satorius) for 120 seconds. Wnt_M1 at 6 or 7 different concentrations went through an association step for 180 seconds and a dissociation step for 180 seconds. The binding kinetics were analyzed using the Octet DataAnalysis 10.0 software package (Satorius). Similarly, biotinylated LRP6 E1E2 (ACROBiosystems) at 400 nM was first immobilized onto streptavidin biosensor tips for 90 seconds, after which association and dissociation were monitored with six serial concentrations of the LRP6-binding protein, following the same protocol used for the Fzd proteins.

### Dual luciferase assay

To assess Wnt signaling activation by the Fzd7 agonist, a dual luciferase assay was performed. HCT116 cells were cultured in DMEM (Sigma) supplemented with 10% fetal bovine serum (Sigma) and 1% penicillin/streptomycin (Cytiva). For reporter assays, cells were seeded and transiently transfected with either TOPFlash or FOPFlash and the Renilla luciferase plasmid pRL-CMV (Promega) using Lipofectamine 3000 (Life Technologies), as previously described ^42^. Cells were treated with recombinant human Wnt3a (R&D Systems), F7_WS_mono or F7_WS_di for 48 h. Firefly and Renilla luciferase activities were measured using the Dual-Luciferase Reporter Assay System (Promega) according to the manufacturer’s instructions.

### Construction of ExpiCHO cells expressing full-length Fzd

Full-length cDNAs for human Fzd receptors (Fzd1-7, 9, 10) were sub-cloned into the pLV lentiviral vector in a Fzd-P2A-muGFP format using standard cloning methods; Fzd8 could not be cloned successfully. Lentiviral particles were produced by transfecting HEK293T cells (T-25 flask, 5 mL DMEM + 10 % FBS, ∼80 % confluency) maintained at 37 °C, 5 % CO_2_. Cells were transfected with 3 µg each of pLV-Fzd and pMD2.G (Addgene #12259) and 1.5 µg each of pRSV-Rev (Addgene #12253) and pMDLg/pRRE (Addgene #12251) using Lipofectamine 3000. Viral supernatant was collected 48 h later, clarified at 300 × g for 2 min, and concentrated to 1 mL with Amicon Ultra-15 filters at 2,000 × g while exchanging into ExpiCHO™ Expression Medium (Gibco). The concentrate was passed through a 0.45 µm PES filter and used to transduce ExpiCHO™ cells (T-25 flask, 5 mL) in the presence of 8 µg/mL Sequabrene. After 48 h, GFP-positive cells were enriched by two rounds of fluorescence-activated cell sorting (FACS) using a CytoFLEX SRT (Beckman Coulter). Approximately 2 × 10⁶ viable cells were pelleted by centrifugation at 300 × g for 2 min, washed once with PBS containing 0.5% bovine serum albumin (PBSF), and resuspended in 400 μL PBSF. After each sorting round, cells were cultured in 5 mL ExpiCHO™ Expression Medium supplemented with antibiotic–antimycotic solution (Thermo Fisher Scientific) for 24 h, followed by maintenance in unsupplemented medium.

### Cell-binding assay by flow cytometry

Each stable cell line, expressing a single Fzd receptor along with muGFP, were washed once with ice-cold PBSF and resuspended at 1×10⁶ cells per 100 μL. Cells were incubated with 100 nM of either Wnt_M1-myc or DRPB_Fz7-myc for 1 hour at 4 °C. After incubation, cells were washed twice and stained with PE-conjugated anti-myc antibody (CST #3739, clone 9B11) for 1 hour at 4 °C. The stained cells were analyzed using a CytoFLEX SRT and data analysis was performed with CytExpert SRT software (Beckman Coulter). Binding was quantified as the percentage of positive cells, gated relative to the untransduced cells treated with each binder.

### Organoid formation from murine Lgr5^+^ intestinal stem cells

Murine intestinal stem cells were embedded in Matrigel (Corning) and cultured in ENR medium composed of Advanced DMEM/F12 (Gibco) supplemented with 1x B27 (Invitrogen), 1x N2 (Invitrogen), 1 mM N-acetylcysteine (Sigma), 250 ng/mL EGF (Peprotech), 100 ng/mL Noggin (IPA), and 10% (v/v) R-spondin1 conditioned medium. Cells were mechanically dissociated and cultured in ENR medium supplemented with F7_WS_mono, F7_WS_di at varying concentrations, or a commercial Wnt surrogate-Fc fusion protein (IPA, Catalog #: N001 - 100 µg) as a positive control. Organoid morphology was evaluated 72 h post-splitting. For confocal microscopic analysis, organoid whole-mount staining was performed, as described ^43^. Briefly, organoids were recovered from Matrigel using Cell recovery solution (Corning) and subsequently fixed in 4% paraformaldehyde. Permeabilization was conducted using 0.3% PBST, followed by blocking with 0.3% PBST containing 1% bovine serum albumin and 2.5% serum. Organoids were sequentially incubated with primary and secondary antibodies for 16 h, and nuclei were counterstained with DAPI. Immunofluorescent images were obtained using an LSM880 confocal microscope after mounting on slide glass.

### *In vivo* hair follicle regeneration assay

All animal procedures were approved by the Institutional Animal Care and Use Committee of KAIST (IACUC No. KA2025-029-v1). Female C57BL/6J mice were obtained from the KAIST Laboratory Animal Resource Center. For intradermal injections, F7_WS_di was freshly dissolved in PBS. Mice (n = 5 per condition) were injected on postnatal day 60. Control mice received the vehicle (PBS) only. A 50 μL volume of F7_WS_di at 0.1 μM, 1 μM, or 4 μM was injected into the dorsal skin once daily on days 1 and 3. Mice were sacrificed on either day 5 or day 14. For proliferation analysis, EdU (25 mg/g body weight in 0.9% NaCl) was administered intraperitoneally 6 h prior to sacrifice. Dorsal skin samples were fixed in 4% paraformaldehyde (PFA) for 1 h at 4 °C, washed with PBS, cryoprotected in 30% sucrose overnight at 4 °C, and embedded in O.C.T. (SAKURA Tissue-Tek) for cryosectioning. For immunofluorescence, 15 μm sections were blocked for 1 h at room temperature with a solution containing 2.5% donkey serum, 1% bovine serum albumin, 1% cold water fish gelatin, and 0.3% Triton X-100 in PBS. Sections were incubated with primary antibody (anti-LEF1, Cell Signaling 2230S) overnight at 4 °C, followed by incubation with secondary antibody and DAPI (Sigma D9542, 1 μg/mL) for 1 h at room temperature. EdU staining was performed according to the manufacturer’s protocol (Click-iT™ EdU Alexa Fluor™ 488 Imaging Kit, ThermoFisher C10339). Images were acquired using a Nikon confocal microscope with a 20× objective and processed using Fiji (ImageJ). Hematoxylin and eosin (H&E) staining was performed according to standard protocols.

### Calculation of pairwise sequence identity

Pairwise sequence identities (PIDs) were calculated for the FZD CRDs. Amino acid sequences were loaded into R using the readAAStringSet function from the Biostrings package. For each sequence pair, a global pairwise alignment was performed using the pairwiseAlignment function from the pwalign package, with the BLOSUM62 substitution matrix applied for amino acid similarity scoring. Following alignment, PID values were calculated using the PID3 method, which defines identity as the percentage of identical positions relative to the length of the shorter sequence in the aligned pair.

## Data availability

The coordinates of the HomoCC structure will be deposited in the Protein Data Bank with the conditions of immediate release upon publication.

## Acknowledgement

This study utilized the Beamline 5C at the Pohang Accelerator Laboratory, Republic of Korea. The research was supported by the National Research Foundation of Korea (Grant Nos. NRF-2021M3A9G8025599, RS-2024-00440039, RS-2023-NR077287, RS-2024-00467046 and RS-2024-00508861) and by the KAIST Convergence Research Institute Operation Program.

## Author contributions

B.-H.O., B.-S.J. and J.L. conceived and designed the research. B.J, H.-C.S. and H.M.K. further refined the concept. J.L. and Y.B. performed the computational design and experimental validation. H.-C.S. and S.J.K. directed and J.M. performed the production and purification of Fzd CRDs. J.C. and Y.K.K. designed and performed the reporter assay. S.Y. and H.Y. designed and performed the *in vivo* hair follicle stem cell activation assay. T.-K.J. and D.-S.L. designed and performed the organoid cultures. B.-S.J. directed and S.A. and H.H. constructed the Fzd-expressing CHO cell lines and performed flow cytometric analysis. B.-H.O., J.L., H.-C.S. and B.-S.J. wrote the original draft. All authors reviewed and accepted the manuscript.

