## Supplementary Figure 1 for "*De Novo* design of a potent Wnt Surrogate specific for the frizzled7 subtype members"

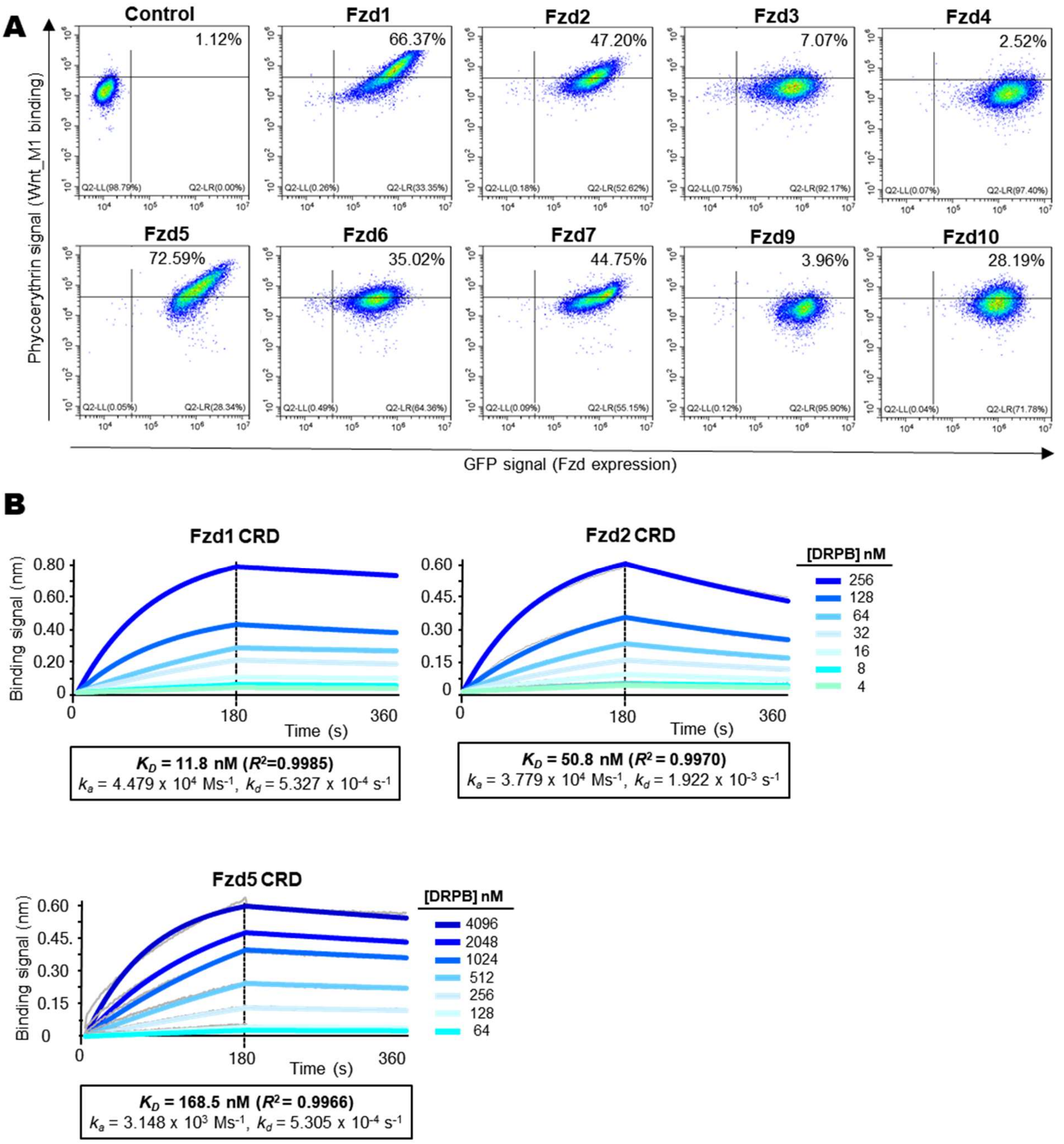


**Supplementary Figure 1. Binding specificity and affinity of DRPB_Fz7.**

**A.** DRPB_Fz7 binding to Fzd-expressing cells. Myc-tagged DRPB_Fz7 (100 nM) was incubated with ExpiCHO cells expressing exogeneous full-length Fzd receptors (as in Figure 2C). Bound ligand was detected by flow cytometry using a PE-conjugated anti-Myc antibody. In the control panel, 1.21% of untransduced ExpiCHO cells exhibited background-level binding signals above the threshold. Considerable binding signals from Fzd5-, Fzd6-, Fzd10-expressing cells are notable.

**B.** Quantification of binding affinities. Biotinylated CRDs of the indicated Fzd subtypes were immobilized to SA-sensor tips and incubated with DRPB_Fz7 at concentrations in a serial two-fold dilution. Binding affinities were estimated using the Octet ForteBio software, with curve fitting performed using a 1:1 steady state model.
