## Supplementary Figure 2 for "*De Novo* design of a potent Wnt Surrogate specific for the frizzled7 subtype members"

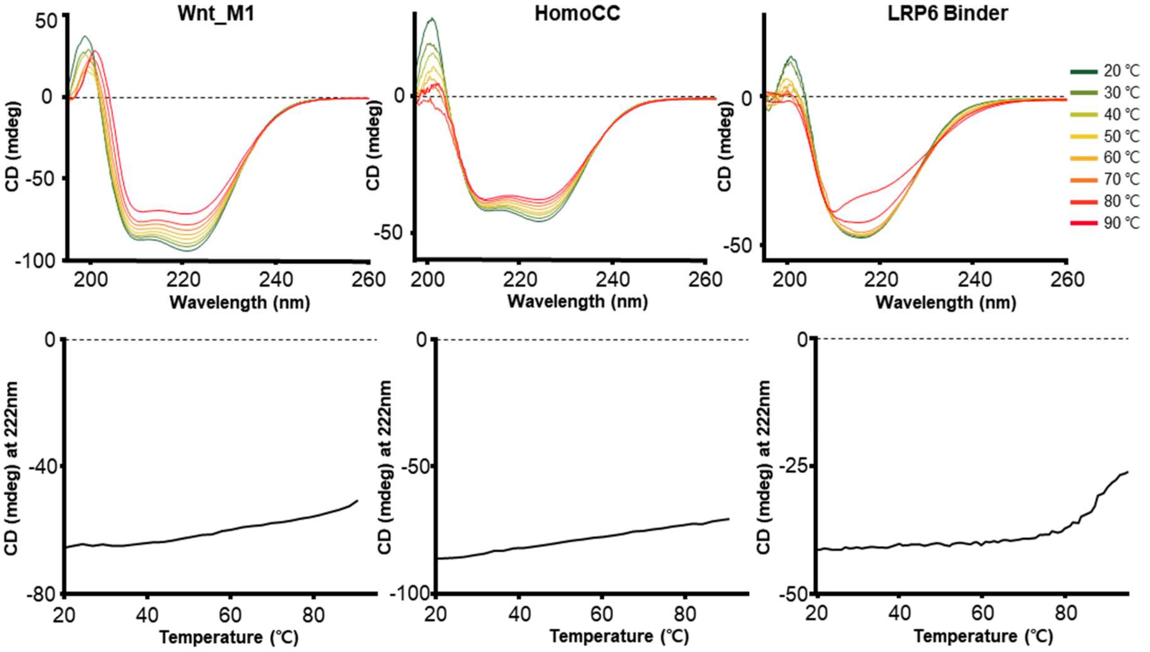


**Supplementary Figure 2. Thermal stability**

Temperature-dependent unfolding of Wnt_M1, HomoCC, and the LRP6 binder was monitored using a JASCO J-1500 CD spectropolarimeter. Wavelength scans from 195 to 260 nm were recorded at the indicated temperatures (*Top*). For melt curve analyses, the CD ellipticity at 206 nm was measured as a function of temperature (*Bottom*).
